# Computer Simulations of the interaction between SARS-CoV-2 spike glycoprotein and different surfaces

**DOI:** 10.1101/2020.07.31.230888

**Authors:** David C. Malaspina, Jordi Faraudo

## Abstract

A prominent feature of coronaviruses is the presence of a large glycoprotein spike protruding from a lipidic membrane. This glycoprotein spike determines the interaction of coronaviruses with the environment and the host. In this paper, we perform all atomic Molecular Dynamics simulations of the interaction between the SARS-CoV-2 trimeric glycoprotein spike and surfaces of materials. We considered a material with high hydrogen bonding capacity (cellulose) and a material capable of strong hydrophobic interactions (graphite). Initially, the spike adsorbs to both surfaces through essentially the same residues belonging to the receptor binding subunit of its three monomers. Adsorption onto cellulose stabilizes in this configuration, with the help of a large number of hydrogen bonds developed between cellulose and the three receptor binding domains (RBD) of the glycoprotein spike. In the case of adsorption onto graphite, the initial adsorption configuration is not stable and the surface induces a substantial deformation of the glycoprotein spike with a large number of adsorbed residues not pertaining to the binding subunits of the spike monomers.

## I. INTRODUCTION

The novel coronavirus SARS-CoV-2 (Figure 1a) emerged in December 2019 as a human pathogen^5^ that causes the COVID-19 disease outbreak that rapidly spread worldwide^6^. This virus belongs to the family of *Coronaviridae* and it is the third documented spillover of an animal coronavirus to humans in only two decades that has resulted in a major epidemic^5^.

**FIG. 1.**
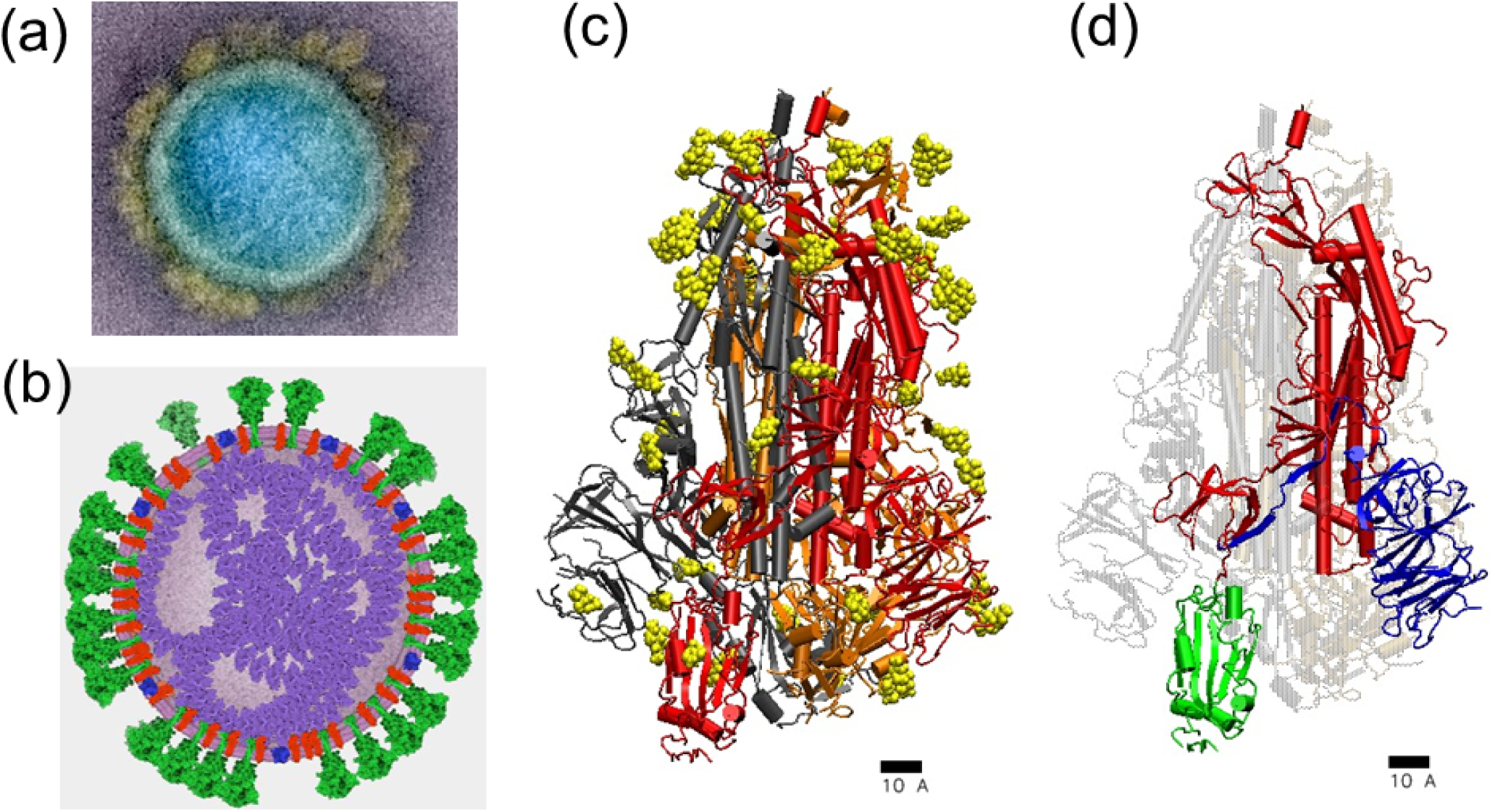
(a) Electron microscopy image of a typical SARS-CoV-2 coronavirus particle, freely distributed by the NIAID’s Rocky Mountain Laboratories (NIAID-RML)^1^, colored to emphasize the virus structure. The spikes protruding from the virus envelope (in yellow color) are clearly visible. Typical diameter ranges from 80 nm to 120 nm. (b) General scheme of a coronavirus indicating their main structural features. We show the nucleocapsid (purple) that packages the viral RNA and the viral envelope. The major ingredients of the envelope are lipids (pink), envelope protein E (in blue), membrane protein M (in red) and the protruding spike glycoproteins (in green). The scheme was made by the authors using CellPAINT^2^. (c) and (d) Snapshots of the atomistic structure of the SARS-CoV-2 trimeric glycoprotein spike available at the Protein Data Bank (PDB:6VSB). The scale corresponds to 1 nm. In (c) the protein is shown in cartoon representation with different colors for each monomer (grey, orange and red). The spike glycosylation is shown in yellow using Van der Waals representation. In (d) the structure of one of the monomers is emphasized. It has a membrane-fusion subunit S2 (in red) and a receptor-binding subunit S1 which has two independent domains (the receptor-binding domain RBD shown in green and the N-terminal domain NTD shown in blue). In this snapshot, the spike was in the prefusion conformation and the RBD shown in green was in its receptor-accessible state (the so-called “up” conformation)^3^. The snapshots were created using VMD^4^.

Coronaviruses are enveloped, single-stranded RNA viruses, with the typical structure shown in Figure 1b. The virus envelope contains lipids and several proteins. These are the so-called envelope (E) and membrane (M) proteins, which play essential roles during virion assembly^7^ and the spike glycoprotein (S) which is responsible for the interaction of a coronavirus particle with a host cell receptor (the ACE2 human receptor, in the case of SARS-CoV-2^8^). The large protruding glycoprotein spikes on the envelope of coronaviruses give a characteristic appearance to this virus family, and give them their name (from “corona”, which is Latin for “crown”).

In an unprecedented effort, the scientific community has been able to rapidly identify not only the nature of the pathogen causing the COVID-19 disease but also most details of its molecular structure with atomistic resolution. For example the identification and full characterization of the virus^9^ was available in February 2020 and the atomistic structure of the spike glycoprotein, shown in Figures 1c,d, was published^3^ as early as in March 2020. At the time of writing, the Protein Data Bank^10^ hosts about ∼300 structures related to the SARS-CoV-2 virus.

This wealth of experimental data has been also augmented with structures obtained from modelling techniques and molecular dynamics (MD) trajectories. For example, in recent works^11,12^ the authors employ modelling software to include in the structure of the virus spike features that are not resolved experimentally (for example, the transmembrane domain) and they use these structures to develop molecular dynamics simulations. Also, the MD trajectories reveal interesting dynamical features^12^. Other theoretical studies consider aspects with direct biomedical implications: investigations of the molecular mechanisms related to the virus infection such as the binding of the virus spike with human receptors^13–15^, identification of targets for vaccine development^16^ and molecular studies related to drug development^17–19^. The exceptionally of the situation also lead to most of the computational groups working in this question to share the structures generated by their models and even full molecular dynamics trajectories, which are being deposited in public repositories such as the COVID-19 Molecular Structure and Therapeutics Hub^20^.

Our aim in this work is to contribute to these computational efforts by considering an important aspect not previously considered in simulation studies, namely the question of the interaction of the SARS-CoV-2 virus with surfaces of materials.

There is substantial evidence that surfaces of materials contaminated by viruses (called fomites in the medical nomenclature) play an important role in human-to-human transmission of many respiratory diseases of viral origin^21–24^, including the particular case of SARS-CoV-2^25^.

Many respiratory viruses are believed to spread from infected people through infected secretions such as saliva or their respiratory droplets, which are expelled when an infected person coughs, sneezes, talks or sings^26^. These respiratory droplets from infected individuals can land on objects, creating fomites (contaminated surfaces)^25^. A recent review of experimental and observational evidence indicates that coronaviruses deposited onto surfaces are able to remain infectious from 2 hours up to 9 days^24^, depending on the surface material and thermodynamic conditions such as humidity and temperature. In the case of SARS-CoV-1 and SARS-CoV-2, evidences from different groups^25^ indicate that viable virus could be detected up to 4 hours on copper, up to 24 hours on cardboard and up to 2-3 days on plastic and stainless steel. This persistence of viable virus onto surfaces is the reason for recommendations of health authorities worldwide on continually disinfecting and cleaning surfaces that are frequently touched.

At the present time, there is a lack of fundamental understanding of interactions between coronavirus and surfaces at the physico-chemical level. We think that such a fundamental knowledge could be very useful in the design and interpretation of experiments involving coronavirus on surfaces and even contribute in the future to the rational design of disinfection measures.

In the case of coronavirus, it seems clear that the presence of the spike coverage in the virus envelope will play an important role in the virus-surface interaction. The spike is not only the most external feature of a coronavirus (see Figure 1) but also a protein which has the ability to interact with other molecules as its main function. Given the fact that the molecular structure and atomistic coordinates of the SARS-CoV-2 virus spike are known^3^, a timely question is to consider the interaction between the spike and surfaces of materials. Starting from the available structure, we perform here atomistic Molecular Dynamics simulations of the SARS-CoV-2 virus spike and surfaces of materials in presence of hydration.

Concerning the materials to be studied, we remark here that previous experimental studies indicated that in general the hydrophobic or hydrophilic nature of the surface plays an important role in the virus-surface interaction^27,28^. Therefore, we will consider here two materials with very different hydrophobic/hydrophilic character: cellulose and graphite. Cellulose is a material which is simultaneously hydrophilic and lipophilic since due to its molecular structure^29^ both hydrogen bonding and the hydrophobic effect play an essential role. In the case of graphite, its surface is strongly lipophilic and mildly hydrophilic^30^, unable to pursue hydrogen bonds and prone to strong hydrophobic interactions. Both materials are widely employed in adsorbents and filters.

Up to the best of our knowledge, this is the first study involving the interaction of the SARS-CoV-2 virus external elements with materials. The results may be also relevant for other coronavirus, since all of them share very similar spike glycoproteins.

## II. SIMULATION RESULTS

### A. Adsorption onto cellulose and graphite

We have performed all-atomic Molecular Dynamics (MD) simulations of a solvated glycoprotein spike near a cellulose and a graphite surface, as shown in Figure 2. As seen in this figure, we have considered the spike inserted inside a large pre-equilibrated water droplet (∼6 × 10^4^ water molecules). The reason for the inclusion of water in the simulation is that it is known that envelope virus (such as SARS-CoV-2) are transferred to surfaces in hydrated conditions (as discussed in the Introduction) and it is also known that the virus needs to be solvated in order to remain viable. The droplet also contains Na^+^ counterions, neutralizing the charge (−23*e*) of the spike (both surfaces are neutral). Full technical details of the models employed and the protocols of the simulations are given in section IV Methods. The process of spike adsorption onto both surfaces is shown in the video provided in the supporting information (SI) and it is illustrated by the snapshots shown in Figure 3. Additional snapshots, showing in more detail the time evolution of the adsorption process during the simulations are provided in the SI. The time evolution of the different quantities characterizing the adsorption process is shown in Figure 4.

**FIG. 2.**
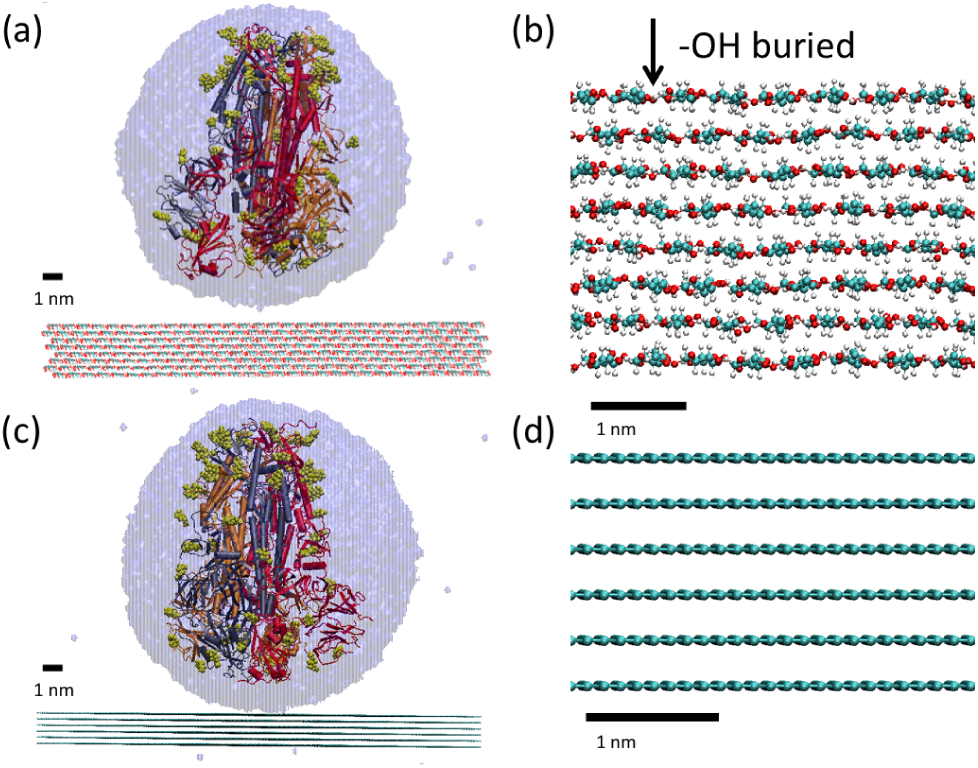
a) Initial configuration for the simulation of the adsorption of a hydrated SARS-CoV-2 spike glycoprotein onto a cellulose surface. The protein is represented as in Figure 1c with its secondary structure with different colors for each monomer of the trimeric protein (red, grey and orange) and the glycosylation in yellow. The solvation sphere is also indicated (transparent blue). b) Detail (side view) of the cellulose surface. Buried -OH groups involved in cellulose-cellulose hydrogen bonds are indicated. c) Initial configuration for the simulationa of hydrated SARS-CoV-2 spike glycoprotein on graphite surface. Color representation is the same as in a). d) Detail (side view) of the graphite surface.

**FIG. 3.**
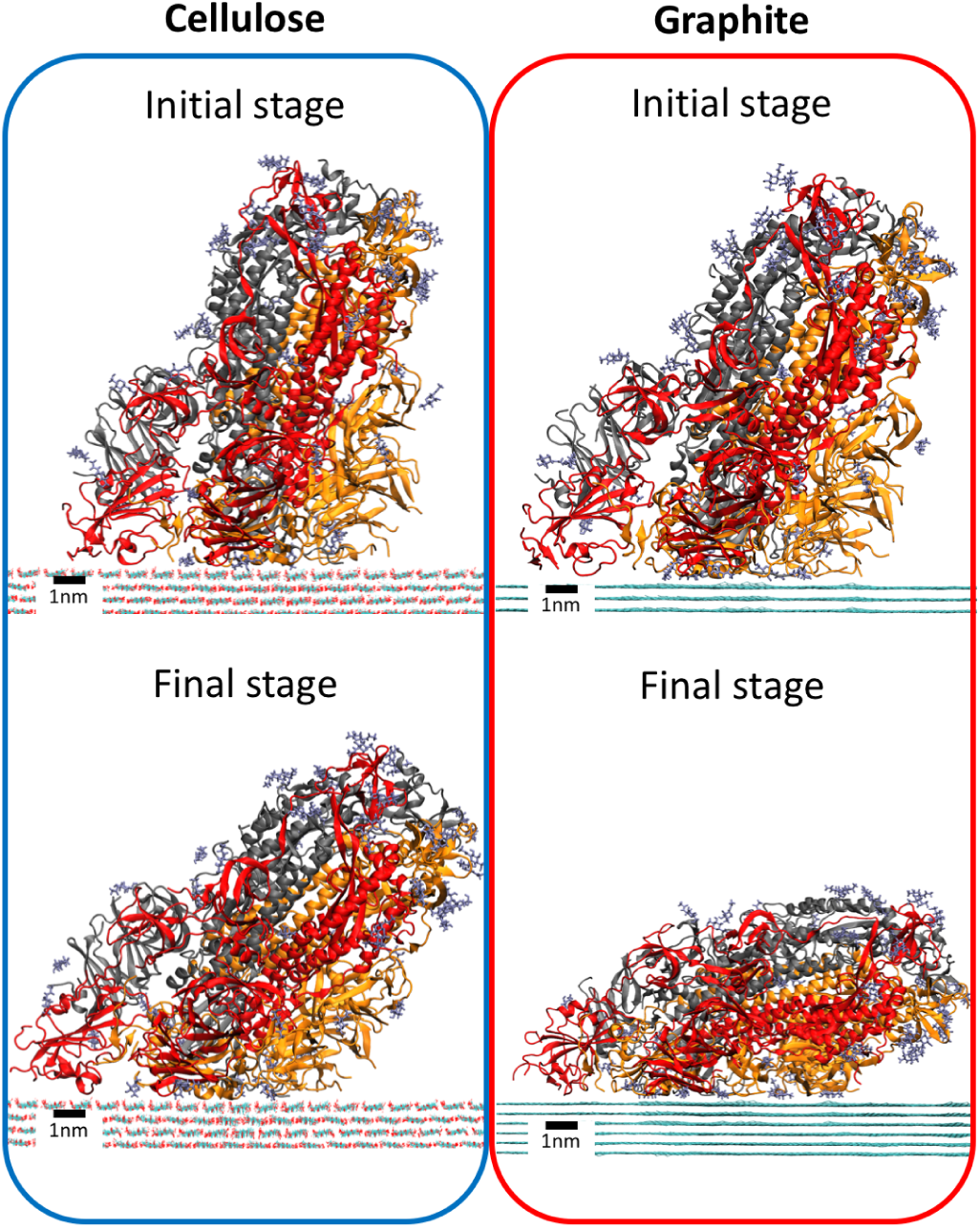
Representative snapshots of SARS-CoV-2 spike glycoprotein during the first and final adsorption stage onto cellulose (left) and graphite (right) surfaces. The initial and final adsorption stage correspond to the time intervals indicated in section IV.Methods. Water molecules were not shown for simplicity (see SI for visualization of the solvation shell). The glycoprotein is shown with cartoon representation and glycans are shown in licorice representation. Each monomer of the trimeric glycoprotein is shown with a different color with the same color code as in Figure 2. The surface is shown with line representation.

**FIG. 4.**
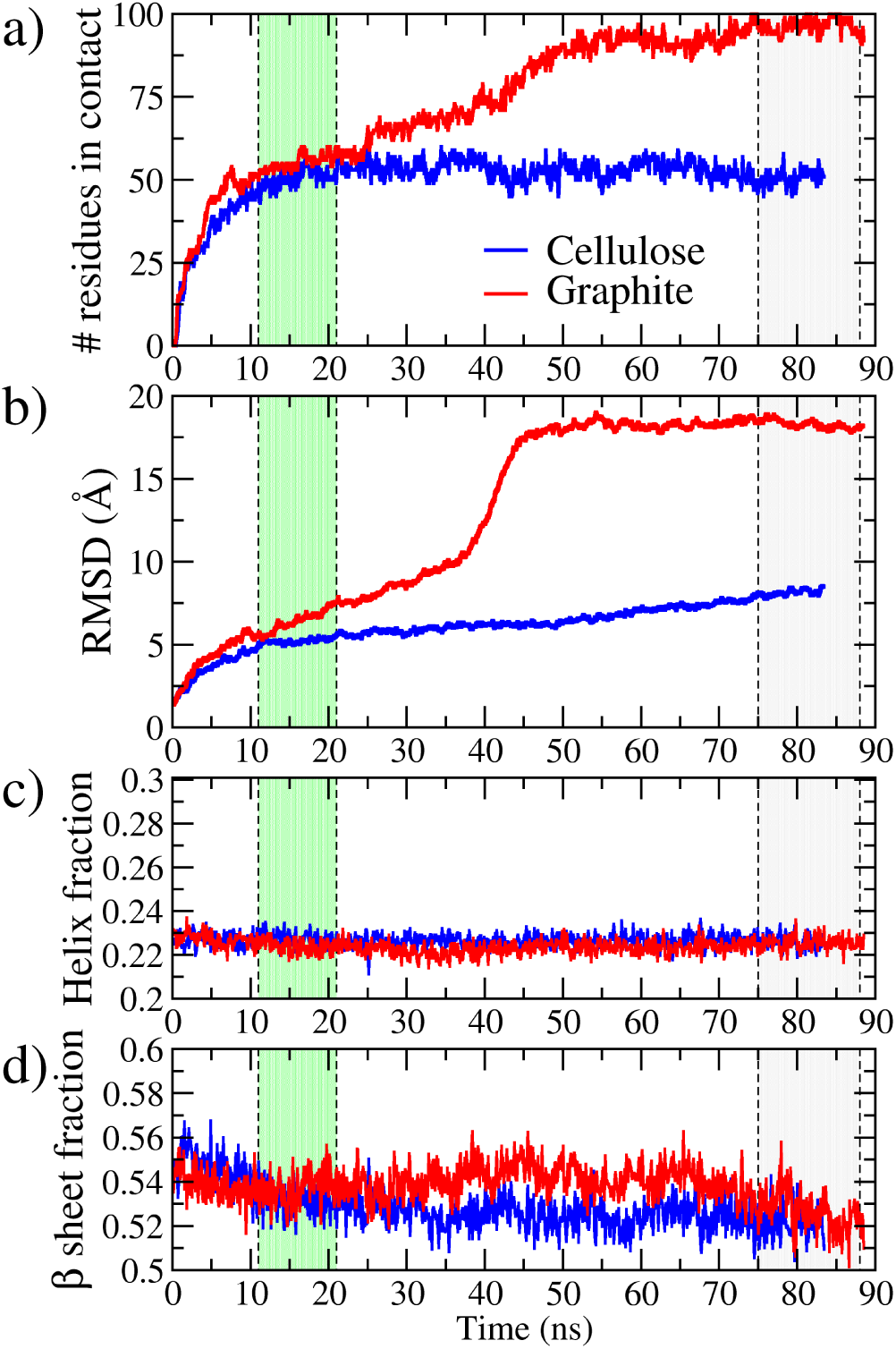
Time evolution of physical quantities in the MD simulations of SARS-CoV-2 spike glycoprotein adsorption onto surfaces. (a) Number of residues in contact with each surface as a function of time. (b) RMSD as function of time. c) Fraction of helix structures in the glycoprotein as function of time. d) Fraction of beta sheet structures in the glycoprotein as a funtion of time. Blue lines correspond to the adsorption on the cellulose surface, while red lines correspond to adsorption on the graphite surface. Yellow and grey areas indicate the approximate location of the initial and final adsorption stage time intervals used in the calculations (see the section IV.Methods for the precise definition).

The simulation results can be briefly summarized as follows. Initially, the spike adsorbs to both surfaces in a similar way (Figure 3), through contact of the receptor binding subunits of the spike with the surface. In the case of cellulose, this configuration is stable and the spike remains essentially in this configuration during all the simulation. In the case of adsorption onto graphite, the initial adsorption configuration is not stable and the surface induces a substantial deformation of the glycoprotein spike.

In order to discuss the results in more detail, it is useful to divide the adsorption process into two different stages, an initial stage corresponding to the contact of the spike with the surface and a final stage reached after structural changes of the spike over the surface.

#### 1. Initial adsorption stage

Full contact between the glycoprotein and the surfaces is established after *t* ∼ 10 ns of simulation in both cases, as indicated by the stabilization of the number of aminoacids in contact with the surface seen in Figure 4a. The number of aminoacids in contact with the surface remains relatively stable for both surfaces (i.e. without abrupt changes) up to *t* ∼ 20 ns, as seen in Figure 4a. We will consider this time interval (with about ∼ 60 aminoacids of the spike in direct contact with the surfaces) as the initial adsorption stage, as highlighted in Figure 4. Illustrative snapshots of this stage are shown in the top panels of Figure 3.

After adsorption (*t* ∼ 10 ns), the RMSD between the adsorbed spike structure in absence of a surface and the adsorbed structure is ∼5 Å for both surfaces (Figure 4b), indicating a small structural change during adsorption. In the case of adsorption onto cellulose, the RMSD remains constant during the initial adsorption stage but it steadily increases with time in the case of graphite. This can be considered as a indication that this adsorbed configuration of the spike onto graphite is not stable, as we will see.

Comparison of the snapshots in Figure 3 and the structure shown in Figure 2d suggests that this initial adsorption of the spike at the surfaces is made through contact between the surfaces and the subunit S1 of each monomer of the spike. This is confirmed by a detailed description of the contact region between the spike and the surface, as shown by the contacts map in Figure 5. As seen in this figure, in both cases (cellulose and graphite), the adsorption involves three RBD and two NTD domains of the receptor-binding subunit S1. In this initial stage, the spike has a similar distribution of contacts with both surfaces although the distribution is more compact in the case of the graphite surface. Therefore, all three monomers are involved in the adsorption process, although in one monomer only the RBD domain is involved and in two monomers both the RBD and NTD domains of the S1 unit are involved. There is only slight contact between the surfaces and glycans or with the S2 subunit of the monomers.

**FIG. 5.**
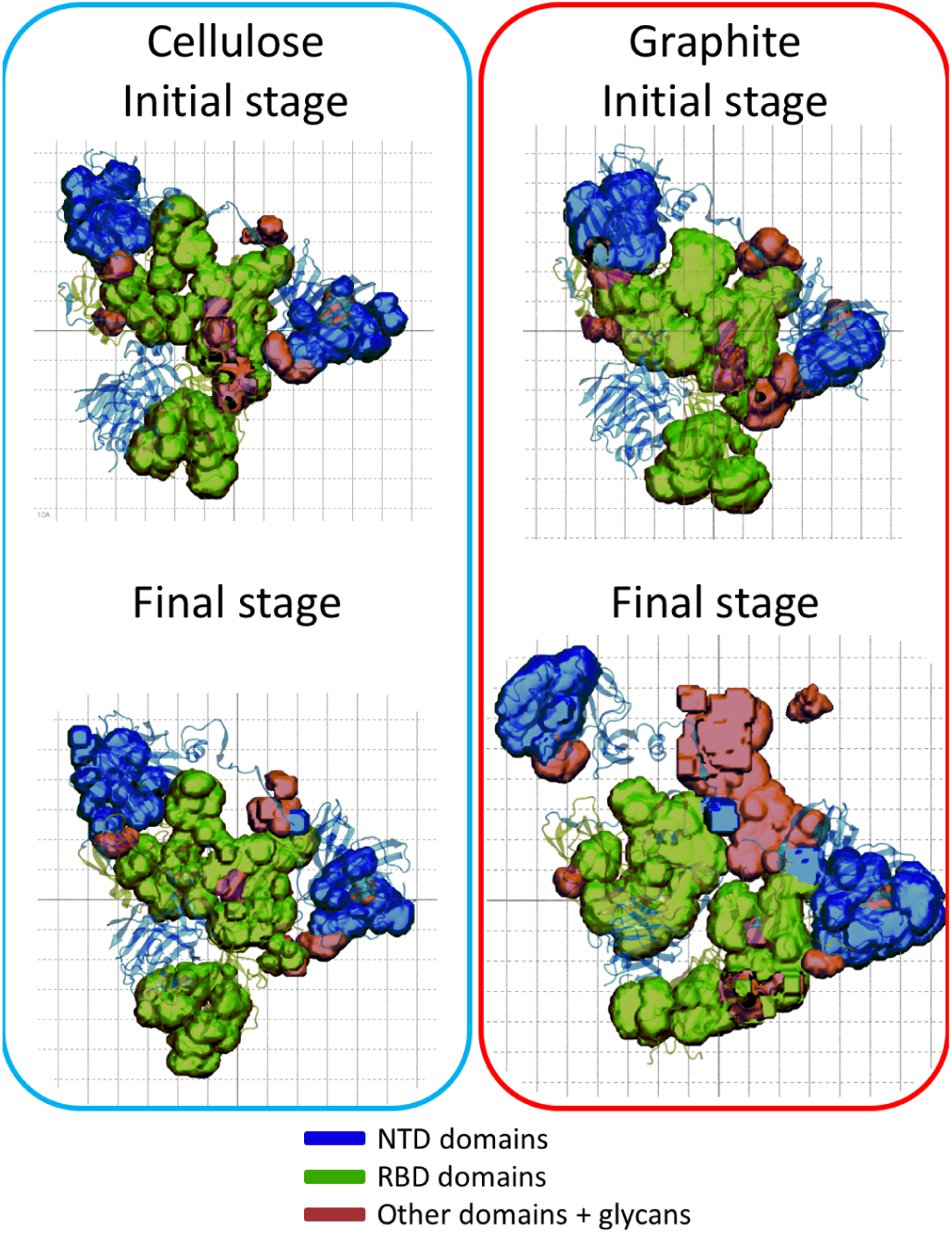
2-D volumetric map of the residues in contact with cellulose (left panels) and graphite (right panels) surfaces during the initial and final stage of adsorption (see section IV.Methods for details). The background grid spacing correspond to 1 nm. Color representation correspond to RBD domain (green), NTD domain (blue) and S2 domain + glycans (red). On top of the volumetric map is the structure of NTD and RBD in transparent cartoon representation.

#### 2. Final adsorption stage

After a similar initial adsorption process, the subsequent evolution reflects (Figures 4a,b) the substantial differences between adsorption onto cellulose and graphite. In the case of the graphite surface, both the number of residues in contact with the surface and the RMSD show substantial evolution with time including abrupt changes (for example at *t* ∼ 40 ns) whereas it shows only minor time evolution in the case of the cellulose surface. At long times (*t* ∼ 75 ns), the spike adsorption onto graphite stabilizes, so we can define a final stage, as indicated in Figure 4, to compare the obtained structures for both surfaces. The most remarkable feature of the final stage is the striking deformation and curvature of the spike towards the graphite surface seen in Figure 3. This deformation is reflected in the large number of contacts of the spike with the graphite surface, which is near 100 residues in contact, as seen in Figure 4a. This deformation of the spike over the graphite surface involve structural changes captured by the time evolution of the RMSD (Figure 4b). The RMSD stabilizes at ∼ 18 Å at the final stage, which implies a substantial structural change induced by graphite.

In the case of the cellulose surface, the number of residues of the glycoprotein in contact with the surface remains approximately constant (∼ 50) between the initial and final adsorption stages. The RMSD remains at ∼ 5 Å up to simulation times of *t* ∼ 50 ns, and after that it increases slowly reaching ∼ 8 Å. This change in RMSD corresponds to a slight deformation of the protein to increase its contact with the surface accompanied by a slight change of the orientation of the main axis towards the cellulose surface (see snapshot in Figure 3).

Figure 5 also shows substantial differences between the surface of contact between the spike and cellulose or graphite, as should be expected from Figure 3. The contacts between the spike and cellulose changed only slightly from the initial to the final stage whereas in the case of graphite the region of contact increased substantially, due to the deformation of the spike discussed above. Figure 5 also shows that in the case of graphite the adsorption involves not only the receptor-binding subunit S1 but also a substantial contact with the membrane-fusion subunit S2. Therefore, the substantial deformation of the spike observed in Figure 3 involves the adsorption of the membrane-fusion subunit S2 at the graphite surface.

Interestingly, neither the adsorption to graphite or cellulose induce changes in the secondary structure of the spike. According to Figures 4b,c there is no significant change in the secondary structure of the spike due to adsorption over surfaces since the percentage of *α*-helix (Figure 4c) and *β* sheets (Figure 4d) structures in the spike remain almost constant.

This could be related to the fact that the spike is known to be rather flexible. In fact, it has been suggested^31^ that the mechanism of biding of the spike of coronaviruses to diverse host cell receptors is based on the flexibility of the spike.

Before entering into a more detailed analysis of the spike-surface interactions, we would like to add a comment about the ions present in the simulation. As we mentioned before, the simulation also contains Na^+^ counterions to neutralize the spike charge. The ions are observed to be mostly condensed at the spike, without being involved in the adsorption process. It is likely that the reason for this observation is that both surfaces are neutral and the spike is strongly charged.

### B. Detailed analysis of protein-surface contacts

In order to obtain a deeper understanding of the adsorption results described in the previous subsection, we have performed a more detailed study of the particular aminoacids involved in the protein-surface interaction. In Figure 6, we show the number of spike aminoacids in contact with cellulose or graphite, classified by aminoacid type, for both the initial and final adsorption stage.

**FIG. 6.**
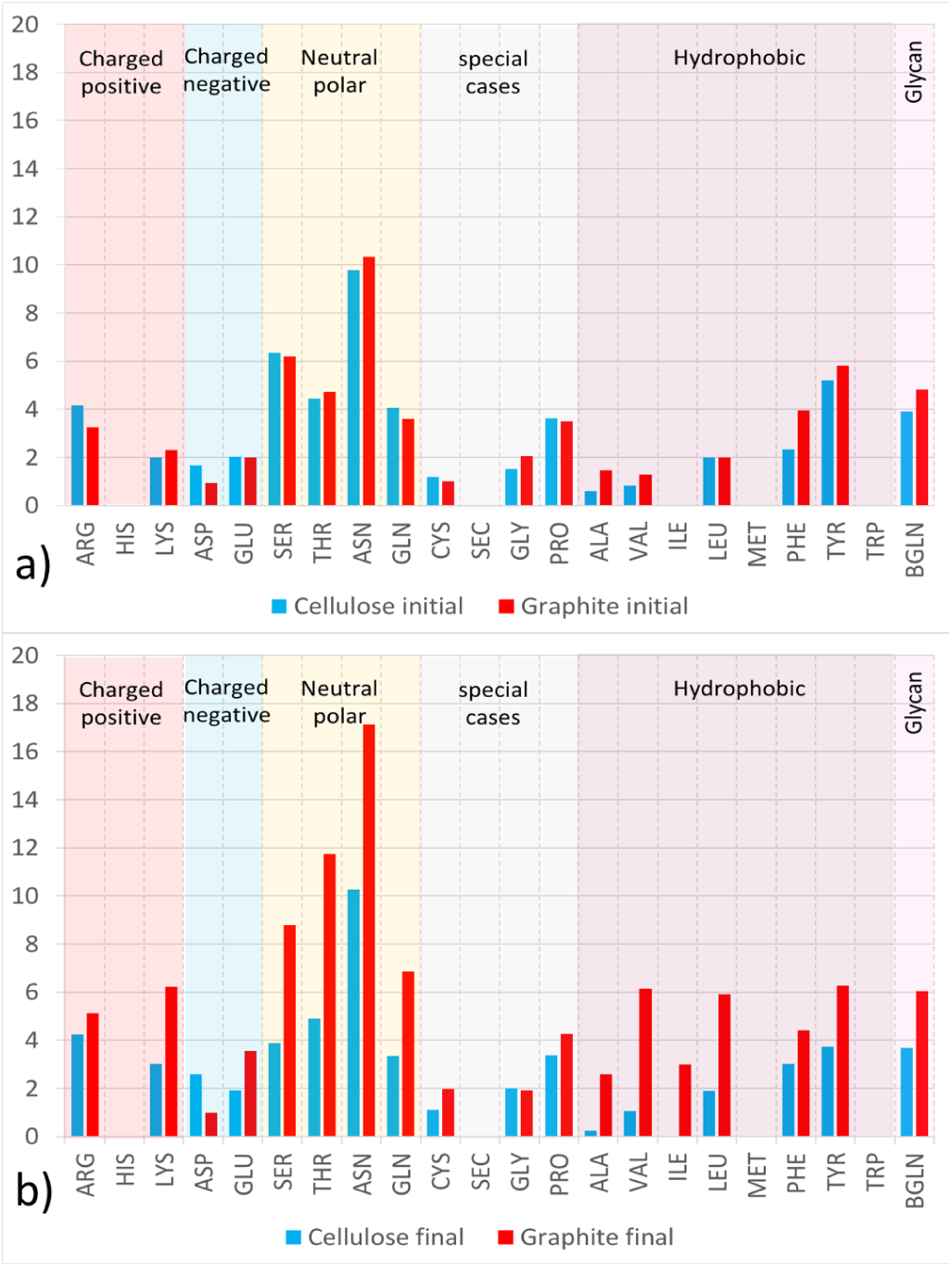
Average number of spike aminoacids (3-letter code) in contact with cellulose (blue) or graphite (red) during (a) the initial stage of adsorption and (b) the final stage of adsorption (see section IV.Methods for details of the calculation). Each aminoacid is also classified into charged (positive or negative), neutral polar, hydrophobic or special cases. Glycans of the spike protein are also included. Standard error bars are too small to be seen at the scale of the figure.

In the initial stage (Figure 6a), we obtain a very similar distribution of residues of the spike in contact with both cellulose and graphite. The only difference is a slight tendency of graphite to favour more contacts with hydrophobic residues. In both cases, there is a substantial contribution from neutral polar aminoacids with ∼24 − 25 contacts (∼ 42-44% of contacts). The most abundant residue in contact with the surface is ASN (Asparagine) with an average of about ∼ 10 contacts (∼17.5% of the total).

Overall, our results in Figures 5 and 6 imply that in the initial adsorption stage, the nature of the surface plays a minor role. In both cases, the spike is able to adsorb to both surfaces through essentially the same aminoacids located in the receptor-binding subunits S1 of the trimeric spike. However, it is possible that the magnitude of the interaction should be different for the different surfaces, given their different character regarding hydrogen bonding and hydrophobic interactions.

In order to compare the magnitude of the spike-surface interactions for both cases, we have performed additional simulations using the steering molecular dynamics technique. In these simulations, the spike is pulled from the surface at constant velocity and the required detachment force is monitored. Our results (reported in Appendix A), indicate that the detachment forces are very similar for both surfaces at the initial adsorption stage. For the faster detachment velocity (5nm/ns) the detachment force for both surfaces is very similar. As we reduce the detachment velocity the difference in the maximum force between graphite and cellulose becomes more important, with graphite requiring a larger force to detach the glycoprotein from the surface (see Figure 8 in Appendix A).

**FIG. 7.**
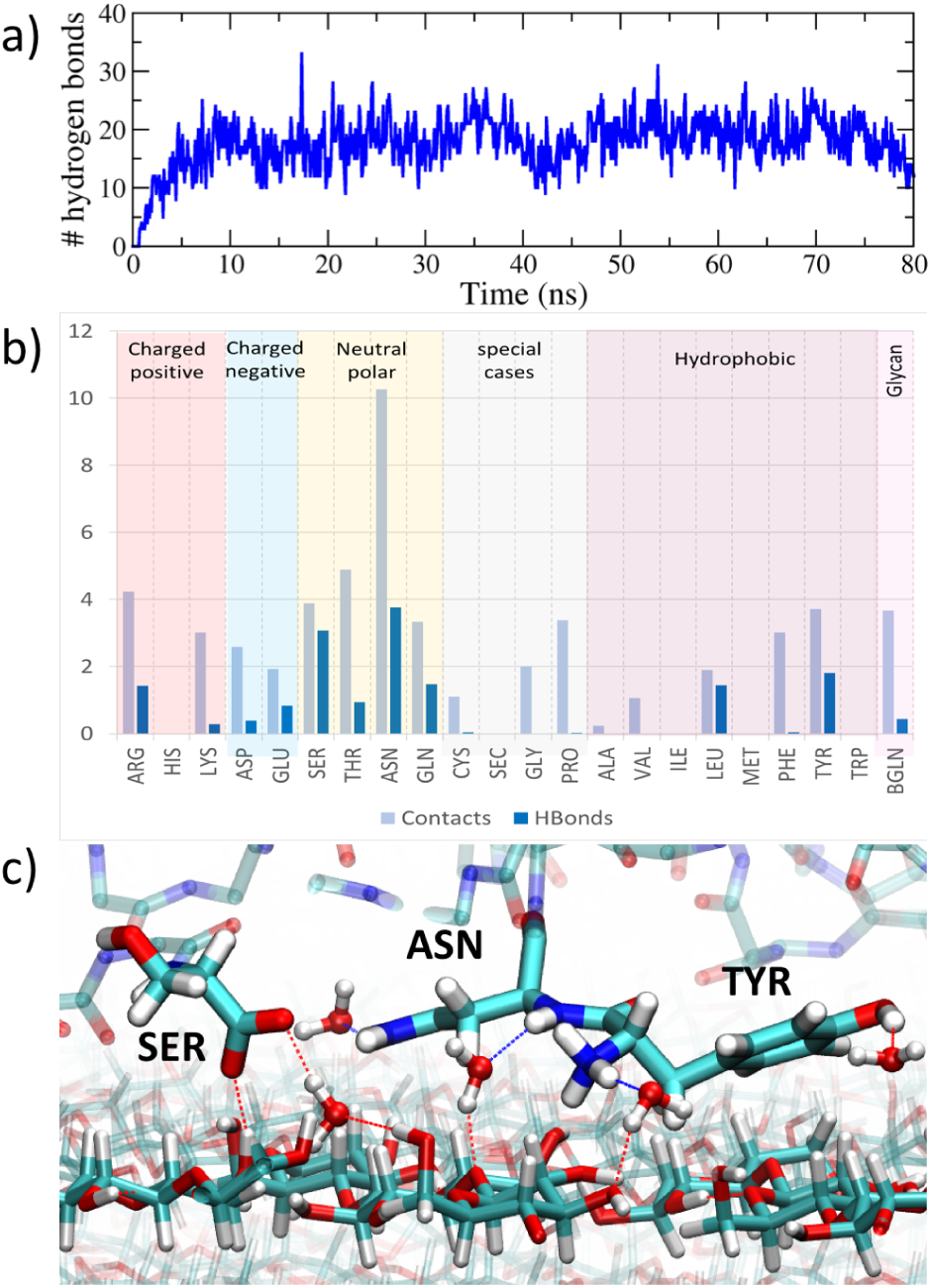
Hydrogen bonds analysis. a) Time evolution of the number of hydrogen bonds between the spike and cellulose. b) Comparison of the distribution of contacts and hydrogen bonds by residues during the final stage of adsorption. c) Simulation snapshot (*t* = 45 ns) with a detail of some spike residues sharing hydrogen bonds with solvation water and cellulose). Highlighted spike residues are SER:443:B, ASN:450:B and TYR:449:B. The dotted lines indicate hydrogen bonds. Both cellulose and aminoacids are shown in licorice representation with CPK colors.

**FIG. 8.**
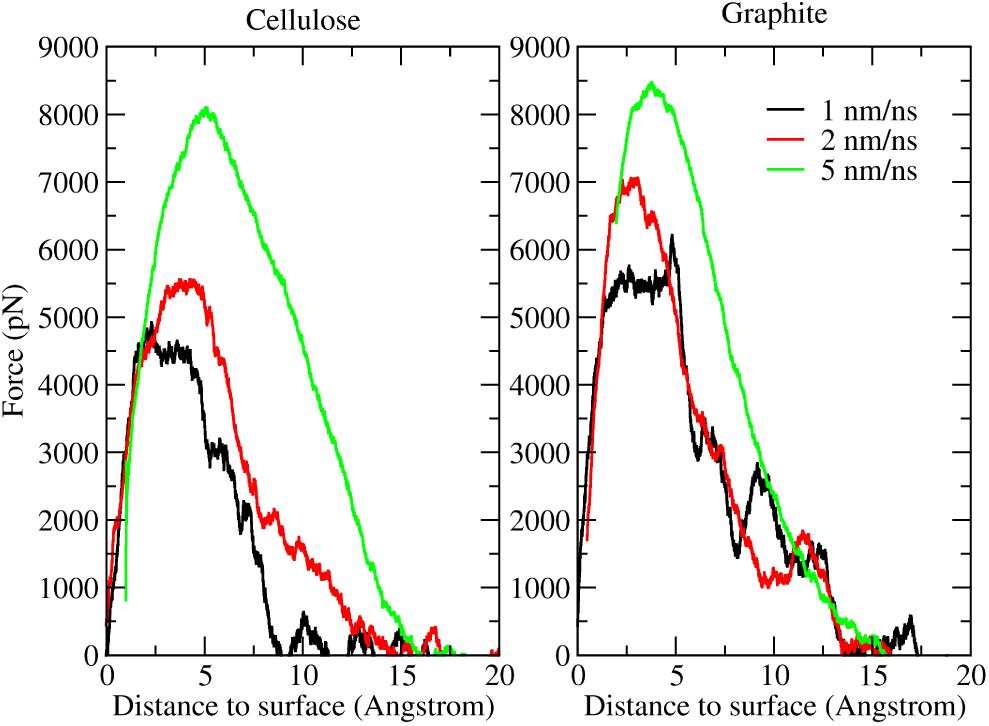
Force as a function of distance, from Stereed Molecular Dynamics (SMD) results, for cellulose (left panel) and graphite (right panel) surface after the initial adsorption stage. Each line correspond to a different pulling velocity: 1 nm/ns is represent with black line, 2 nm/ns with a red line and 5 nm/ns with a green line.

This higher detachment force for the graphite surface could be related to the density of contacts as shown in Figure 5 at the initial adsorption stage. As seen in that figure, the graphite tends to form a more dense contact surface with the spike glycoprotein, which may require a larger force to detach from the surface. In any case, this comparison between detachment forces should be considered with caution, since these SMD simulations are noisy, experiencing substantial fluctuations.

The results for the analysis of the aminoacids involved in the protein-surface interaction during the final stage (Figure 6b) reflect the different evolution for the adsorption of the spike on cellulose or on graphite, in line with the results discussed in the previous subsection. In the case of graphite, the differences between the initial stage and the final stage in Figure 6 are obviously due to the deformation of the spike after adsorption described in the previous subsection. The total number of contacts of the spike with graphite increased from an average of ∼ 59.2 contacts in the initial stage to ∼ 102.8 in the final stage.

Again, the most abundant residue in contact with the graphite surface is ASN (asparagine) with an average of about ∼ 17 contacts corresponding to a prominent peak in Figure 6b. This result is consistent with previous simulations that indicated a strong affinity of asparagine with carbon aromatic rings^32^. The final stage of adsorption at the graphite surface is dominated by contacts with asparagine, threonine, serine and glutamine neutral polar aminoacids but there is also a substantial number of contacts with hydrophobic aminoacids such as tyrosine, valine or leucine and with the glycans covering the lateral regions of the spike.

In the case of cellulose, the number of contacts remains nearly the same (only a very slight decrease in the total number of contacts, from a total of 55.6 in the initial stage to 54.2 in the final stage). Comparison between Figure 6a and Figure 6b shows very minor changes. In the final stage there are slightly more contacts with charged aminoacids and less contacts with hydrophobic aminoacids than those obtained in the initial stage. Therefore, the small changes observed in the previous subsection (both in the RMSD and the map of contacts, Figures 4b and 5) can be attributed to a rearrangement of the protein at the surface to increase interactions with hydrophilic aminoacids and reduce the contacts with hydrophobic aminoacids, without significantly changing the number of contacts. In any case, Figure 6 shows a wide variety of residues with different chemical affinity in contact with cellulose. Again the peak in the case of asparagine is the most noticeable feature for the case of cellulose in Figure 6b, with ∼ 10 contacts (which corresponds to ∼19% of contacts).

### C. Detailed analysis of protein-cellulose hydrogen bonds

Overall, our results indicate that in the case of cellulose the spike is immobilized after adsorption, experiencing only minor changes during the adsorption process. This effect could be due to some sort of stabilizing interaction, that anchors the aminoacids of the receptor binding domain (RBD) after touching the surface.

Since the surface of cellulose has a large hydrogen bonding ability, this interaction could be responsible for the observed stabilization. In order to check this possibility, we have analyzed in detail the presence of hydrogen bonds between the spike and the cellulose surface (Figure 7). In Figure 7a, we show the number of direct hydrogen bonds between the spike and the cellulose surface. Similar to what is observed in Figures 4a and 4b with the number of contacts and the RMSD, the number of hydrogen bonds stabilizes after ∼ 10 ns of simulation time and fluctuates around an average of ∼18 hydrogen bonds total during the rest of the simulation. Further identification of these hydrogen bonds reveals that they are mostly located on the RBD domain of the glycoprotein, constituting almost ∼ 75% of the total hydrogen bonds.

In Figure 7b, we report the number of hydrogen bonds for each aminoacid type together with the number of aminoacid-spike contacts. In the case of the neutral polar asparagine and serine, a significant number of the aminoacid-cellulose contacts involve hydrogen bonding. It is also interesting to note a significant number of hydrogen bonds with cellulose from the aminoacids of hydrophobic character leucine (LEU) and tyrosine (TYR) that have also the possibility of hydrogen bonding. Probably the amphiphilic character of cellulose^29^ tends to enhance the interaction with these aminoacids.

Hydrogen bonding between the spike and cellulose is more complex than simply due to direct cellulose-spike hydorgen bonds. A closer look to the formation of hydrogen bonds between the spike and cellulose reveals the existence of hydration water molecules that share hydrogen bonds with cellulose and aminoacids. In Figure 7c we can observe that three of the aminoacids with larger contributions in the number of hydrogen bonds in Figure 7b (ASN, SER and TYR) also form bridging hydrogen bonds with surrounding water, which makes hydrogen bonds with both the aminoacid and the cellulose surface.

Overall, this complex hydrogen bond network mainly located at the interface of the RBD domain tends to stabilize the spike glycoprotein on the cellulose surface and is possibly responsible for the differences in deformation observed between cellulose and graphite surfaces observed in Figure 3.

## III. CONCLUSIONS

In this work we presented molecular dynamics simulation of the SARS-CoV-2 spike glycoprotein interacting with two different surfaces: cellulose and graphite. The choice of these surfaces was made in order to compare two different materials with very different properties. Cellulose is a complex molecular material with amphiphilic properties and a high quantity of hydrogen bonds donors and receptors. Previous works (see for example ref^29^ and references therein) demonstrated the capacity of cellulose to bind proteins by both hydrogen bonding and hydrophobic interactions. On the contrary, graphite is a crystalline hydrophobic material with no hydrogen bond capability. It is also known to be able to bind peptides and proteins via hydrophobic interactions (see for example^32^ or the discussion in Ref^33^).

Our simulation results can be summarized as follows:

Initially, the spike adsorbs to both surfaces through essentially the same residues belonging to the receptor binding subunit of its three monomers (in particular, involving all three receptor-binding domains (RBD) and two N-terminal domain (NTD)). From this point the adsorption on each surface dramatically differs.

Adsorption onto cellulose stabilizes in the initial adsorption configuration with the help of a large number of hydrogen bonds developed between cellulose and the three receptor binding domains (RBD) of the glycoprotein spike. This adsorbed configuration also includes shared hydration water between the spike and cellulose. In the case of adsorption onto graphite, the initial adsorption configuration is not stable and the surface induces a substantial deformation of the glycoprotein spike with a large number of adsorbed residues not pertaining to the binding subunits of the spike monomers.

It is interesting to note that our results are in line with previous MD results of other proteins at these surfaces. Cellulose tends to adsorb proteins in stable configurations without structural changes^29^ whereas the interaction with graphite induce substantial structural effects on adsorbed proteins^33^.

Concerning the possible practical implications of these results, obviously we need to remark that the present study is a simplification, since it ignores important effects such as the process of approach of the full virus to the surface, which is dominated by long range forces. Nonetheless, this represents the final stage of the adhesion of a virus with a surface, in which the most external element (the spike) interacts with the surface and it provides a reasonable approximation to the affinity between the virus particle and a given surface. With all these precautions in mind, we can say that our results suggest that interactions with cellulose will tend to maintain the integrity of the hydrated SARS-CoV-2 virus spike. Also, interactions with graphite deform the spike and may potentially help to inactivate the infectious potential of the spike glycoproteins interacting with the surface. As a recommendation for future experimental investigations, it will be of great interest to investigate the viability of the virus over carbon surfaces, in particular given the importance of these materials for filtration applications.

Our study has to be considered a first step in the understanding of the molecular interactions between the SARS-CoV-2 virus and surfaces. Of course, our study has many limitations and further work is necessary in order to understand many relevant factors that are beyond the scope of this paper. One obvious limitation is that in our simulations we considered only the (hydrated) spike glycoprotein of the SARS-CoV-2 virus but not the virus a whole.

The molecular scale analysis of the virus-surface interaction is only one of the relevant aspects that need to be considered in order to understand the interaction between this SARS-CoV-2 virus and materials. Other factors operating at larger length scales need also to be considered. For example, experimental studies for other viruses show evidence that porosity and nanostructuration of the surfaces at scales of the order of the virus size also have an impact^28^.

Also, modelling of the respiratory droplets embedding the virus (which contain mucosal biopolymers, lipids and salts^34^) and how these droplets interact with materials and textiles is of the highest interest. A simulation study of these factors will require the use of mesoscale models, which may be build from relevant experimental data -which is still unavailable- or eventually from the results of atomistic molecular modelling, as has been done recently for mesoscale simulations of a full influenza virus^35^.

## IV. METHODS

### A. Simulation models and forcefields

All Molecular dynamics (MD) simulations reported in this paper were performed using NAMD 2.13 software^36^. The preparation of the simulation models and most of the analysis were made using Visual Molecular Dynamics (VMD) software^4^. The force field employed in the simulations is the CHARMM36 force field which includes parametrization of carbohydrate derivatives, polysaccharides and carbohydrate–Protein interactions^37^. This forcefield is therefore appropriate for describing both the spike glycoprotein and all materials considered in the paper. The water model used in our simulations was the TIP3P model included in CHARMM36.

The atomic coordinates for the SARS-CoV-2 spike glycoprotein structure were obtained from a cryo-EM structure^3^ solved at 3.46 Å average resolution (PDB ID: 6VSB). This structure contains S1 and S2 spike subunits (with one RBD domain in “up” conformation) and a glycosylation pattern characterized by N-Acetyl-D-Glucosamine (NAG) residues, as shown in Figures 2c and 2d. The only modification made to this initial structure was the addition with VMD of missing hydrogen atoms and connecting links between the protein aminoacids and the NAG residues. The obtained structure contains 46,708 atoms and its total charge (assuming pH 7) is −23e.

It should be noted that the glycosylation pattern present in this structure only includes glycans in close proximity to the protein due to lack of further information on the resolved structure of the spike. We are aware of ongoing work on the development of more accurate models of the spike in order to include details not resolved in the available structures^12,16^ such as improved models of the glycosylation. We think that these details, which are essential in questions such as recognition of the spike by the immune system or its interaction with specific receptors will not be essential in the study of the interaction of the spike with extended surfaces. In any case, developments on improved spike models should be carefully considered in future simulations of the virus interactions with materials.

The spike structure was solvated using VMD with an spherical solvation shell in order to maintain its hydrated functional state. The number of TIP3P water molecules added to solvate the glycoprotein was 60,642. We also added 23 Na^+^ counterions to neutralize the charge of the spike. The system made by the hydrated spike with counterions has a total of 228,657 atoms.

The structures of the surfaces were built as follows. The cellulose structure was built using the Cellulose builder toolkit^38^ from a cut of the crystallographic plane (100) from cellulose I*β* crystal structure as in our previous work^29^. We selected the (100) cellulose surface because it is the structurally simplest and smoothest surface that can be generated from cutting the I*β* cellulose crystal structure (see for example Figure 2 in Ref^29^). In any case, our previous studies^29^ show that the different surfaces of cellulose have similar wetting properties and similar hydrogen bonding capacity. An interesting feature of the (100) cellulose surface is that it has “buried” -OH groups involved in cellulose-cellulose hydrogen bonds that can be broken to generate hydrogen bonds of cellulose with adsorbing molecules (see for example Fig.3 in Ref^29^). The generated cellulose structure has a surface with dimensions of 26.1 nm × 25.08 nm and a thickness of 3.18 nm (8 molecular layers) as seen in Figure 2b. The cellulose structure has 252,000 atoms and the full simulation box with the hydrated spike, Na^+^ counterions and cellulose has 480,633 atoms.

The graphite structure was build using the inorganic builder plugin of VMD^4^ by replicating the unit cell 100 times in *a*, 100 times in *b* and 3 times in *c* direction. A detail of the surface can be observed in Figure 2c. Since graphite has a hexagonal crystal structure, we used also periodic boundary conditions with the same geometry, with simulation box vectors (in nm) *a*=(12.28,-21.27,0.00), *b*=(12.28,21.27,0.00), *c*=(0.00,0.00,40.0). The graphite structure has 120,000 atoms and the full simulation box (hydrated spike, counterions and surface) contains 348,654 atoms.

We recall here that all surfaces considered in our simulations are neutral.

### B. Molecular Dynamics simulations protocols

The protocol followed in all simulations includes an initial minimization, equilibration and production runs. In all simulations Newton’s equations of motion were integrated with a 2 fs time step and electrostatic interactions were updated every 4 fs. All bonds between heavy atoms and hydrogen atoms were keep rigid. All simulations were performed in the NVT ensemble with a Langevin thermostat set at 298 K and a damping coefficient of 1 ps^−1^. We employed periodic boundary conditions in all directions. Lennard-Jones interactions were computed with a cutoff of 1.2 nm and a switching function starting at 1.0 nm. Electrostatic interactions were computed using Particle Mesh Ewald (PME) algorithm using a real space cutoff set at 1.2 nm and a PME grid of 1.0 Å.

We performed three different MD simulations. First, we performed a preliminary simulation (19 ns) of the solvated spike at 298K. Employing the results of the preliminary simulation as starting configuration, we have performed MD simulations of the protein spike adsorption onto cellulose and graphite. The equilibrated spike inside a water droplet was positioned at 2 Å away from the surface, as shown in 2. The simulation trajectory was run for 83.4 ns in the case of adsorption onto cellulose and 88.5 ns in the case of graphite.

### C. Analysis of results

The snapshots and movies of the simulations were made using Visual Molecular Dynamics (VMD) software^4^. The different analysis were made using VMD tools and appropriate scripts as follows.

As discussed in the main paper, for convenience in the analysis we introduce an initial adsorption stage and a the final adsorption stage. In the calculations of averaged quantities, the exact definition of these stages is as follows. We define the initial adsorption stage as the time interval between 12.55-19.44 ns for simulations of adsorption over the cellulose surface and between 11.0-21.0 ns for the case of graphite surface. Similarly, we define the final adsorption stage as the time interval between 74.5-83.4 ns for the simulation with cellulose and between 78.5-88.5 ns for the simulation with graphite. Note that this choice of time intervals is related to simplicity in data handling and the time intervals shown in Figure 4 are not exact (since the exact definitions slightly differ for each surface) but approximate for illustrative purposes.

The number of aminoacids in contact with each surface (Figure 4a) was computed considering that a contact between aminoacids and surface occurs when at least one atom of the aminoacid is found at a distance smaller than 3.5 Å from any surface atom. In order to co count the number of aminoacids at each time timestep we employed a TCL script running on VMD implementing the distance requirement described above. The distribution of residues in contact with the surfaces (Figure 6) was calculated over the initial and final adsorption stage with a similar TCL script, averaging over the intervals defined above. The 2-D contact map (Figure 6) was calculated using VMD Volmap tool for residues at distance of less than 3.5 Å from surface atoms. The root mean squared deviation (RMSD) reported in Figure 4b was computed between each instantaneous structure and the initial structure using the RMSD trajectory tool implemented in VMD^4^. The analysis of secondary structure as function of time in Figures 4c and 4d was made using the timeline tool in VMD, which uses the STRIDE algorithm^39^ to calculate the fraction of different secondary structure components. Hydrogen bonds (Figure 7) were computed using VMD. We used an acceptor-donor distance cutoff of 3.5 Å and acceptor-hydrogen-donor angle cutoff of 30 degrees.

## ACKNOWLEDGMENTS

This work was supported by the Spanish Ministry of Science and Innovation through grant RTI2018-096273-B-I00 and the “Severo Ochoa” Grant SEV-2015-0496 for Centres of Excellence in R&D awarded to ICMAB. We thank the CESGA supercomputing center for computer time and technical support at the Finisterrae supercomputer. D. C. Malaspina is supported by the European Union Horizon 2020 research and innovation programme under Marie Sklodowska-Curie COFUND grant agreement No. 6655919.

## Appendix A: Protein detachment by Steered Molecular Dynamics

The detachment force of the spike at cellulose and graphite surfaces adsorbed at the initial adsorption stage (Figure 3) was calculated using the Steered Molecular Dynamics technique^40^ (SMD) as implemented in NAMD. The SMD simulations were conducted starting from the configuration obtained in the MD simulations at 21.0 ns for adsorption onto the cellulose surface and 19.44 ns for the graphite case. The spike glycoprotein was pulled from the center of mass of the residues located at less than 4nm from the surface, this roughly correspond to the RBD and NTD domains. The parameters for the SMD simulation are the same as previous simulations with the addition of a forcing to the spike (force constant 2 × 10^4^ kcal/mol/Å^2^) ensuring a constant velocity of pulling that was set to 1 nm/ns, 2 nm/ns and 5 nm/ns. According to these velocities the simulation time was selected in order to obtain a separation of at least 2 nm between the spike and the surface.

As a result, in SMD we obtain force-separation curves corresponding to each pulling velocity. These forces as a function of spike-surface distance obtained in the SMD simulations were rather noisy (as usual in SMD simulations) so they were smoothed with a running average.

We should keep in mind that the obtained forces from the SMD simulations correspond to nonequilibrium processes in which the motion of the spike will experience a viscous drag (which depends on velocity) in addition to the adhesion force. This viscous resistance can be identified by noting that the force should decay to zero as the protein separates from the surface. In the SMD simulations, we observe a decay of the force with distance to an approximately constant value. This value can be taken as an approximation to the viscous resistance. Therefore, in order to remove the effect of viscous drag and extract the adhesion force, we have shifted the force versus distance curves obtained in SMD so that they decay to zero force at large spike-surface separations. The values of the estimated viscous drag depend on the spike velocity in the SMD simulations. For the simulations with the cellulose surface they were 4,933 pN, 6598 pN and 11,150 pN for SMD simulations of velocities of 1nm/ns, 2nm/ns and 5 nm/ns respectively. Similarly, the values in the case of simulations with the graphite surface for SMD simulations with velocities 1nm/ns, 2nm/ns and 5 nm/ns were 4,390 pN, 7,489 pN and 11,797 pN respectively. The force versus distance curves obtained after this process were shown in Figure 8.

For the faster detachment velocity (5nm/ns) maximum force in both surfaces is approximately similar, been ∼8100 pN for cellulose surface and ∼8500 pN for the graphite surface. As we reduce the detachment velocity the difference in the maximum force between graphite and cellulose becomes more important, with graphite requiring a larger force to detach the glycoprotein from the surface. At the lower detachment velocity (1 nm/ns) the maximum force for cellulose surface is ∼4900 pN and ∼6200 pN for the graphite surface.

## Notes

### Competing Interest Statement

The authors have declared no competing interest.

### Summary of Updates

Corrected Figure 7c (visualization of hydrogen bonds and several aminoacids was confusing). Several references were updated. Minor changes in the main text. Updated Conclusions section. Added supplemental images with time evolution of the simulations.

