## Supplemental Information for "Computer Simulations of the interaction between SARS-CoV-2 spike glycoprotein and different surfaces"

(Dated: 25 September 2020)

**The following article has been submitted to the journal *Biointerphases*. After it is published, it will be found at https://avs.scitation.org/journal/bip**

### I. ADDITIONAL SIMULATION SNAPSHOTS AT DIFFERENT TIMES

Here we provide additional simulation snapshots for the MD simulations reported in the main paper. Figure S1 corresponds to the case of adsorption onto cellulose and Figure S2 corresponds to the case of adsorption onto graphite (see details in the main text). The snapshots were taken at different times so that the process of protein approach to the surface and initial and final adsorption stages can be visualized. In the snapshots we include not only the surface and the spike protein but also we indicate the approximate location of the solvation shell that covers the surfaces and the protein.

a)Electronic mail:

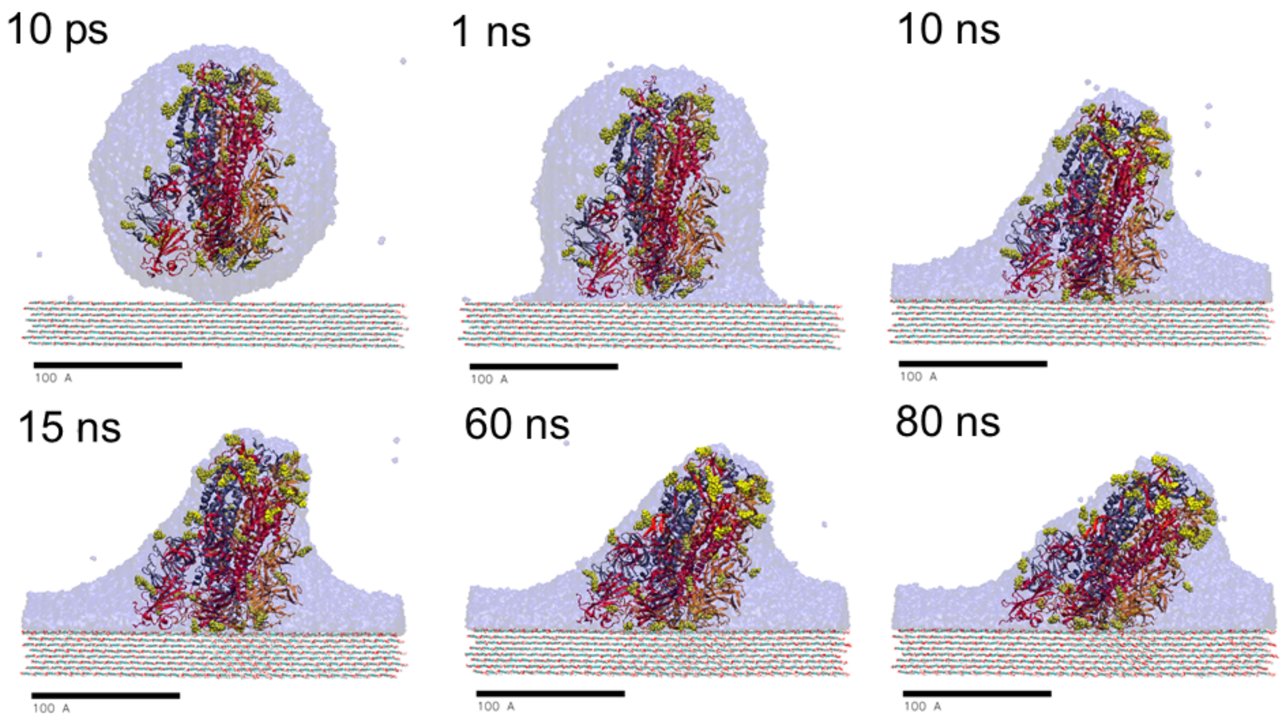


FIG. 1. Snapshots taken at different times of the MD simulation of the adsorption of a hydrated SARS-CoV-2 spike glycoprotein onto a cellulose surface. The protein is represented as in the main paper with its secondary structure with different colors for each monomer of the trimeric protein (red, grey and orange) and the glycosylation in yellow. The volume occupied by the solvation water is also indicated (transparent blue). The surface is represented with lines. The scale bar corresponds to 100 Å.

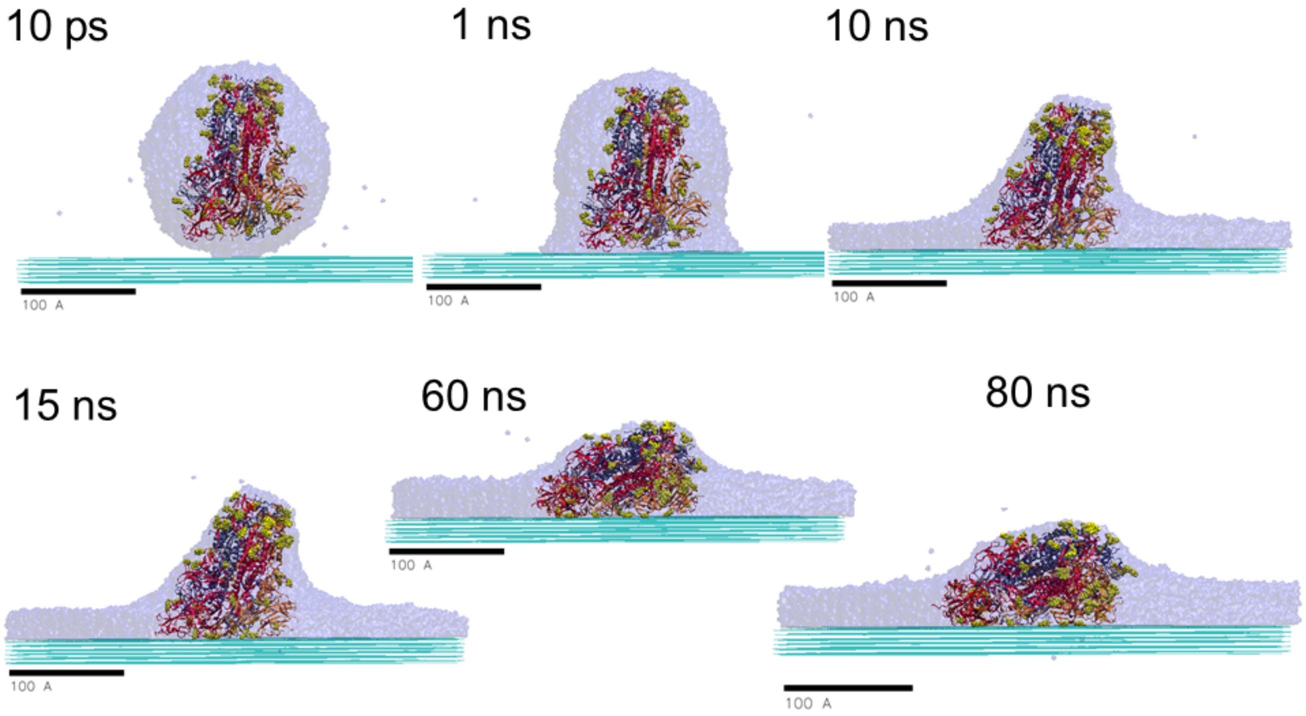


FIG. 2. Snapshots taken at different times of the MD simulation of the adsorption of a hydrated SARS-CoV-2 spike glycoprotein onto a graphite surface. The color codes and representations are the same as in the previous figure.
